# Mood variability during adolescent development and its relation to sleep and brain development

**DOI:** 10.1101/2022.08.23.505008

**Authors:** Yara J. Toenders, Renske van der Cruijsen, Jana Runze, Suzanne van de Groep, Lara Wierenga, Eveline A. Crone

**Affiliations:** Developmental and Educational Psychology, Leiden University, the Netherlands; Leiden Institute for Brain and Cognition, Leiden University, The Netherlands; Department of Psychology, Education and Child Studies, Erasmus University Rotterdam, the Netherlands; Clinical Child and Family Studies, VU University Amsterdam, the Netherlands

**Keywords:** adolescence, mood variability, brain structure, sleep, actigraphy, anxiety, depression

## Abstract

Mood swings, or mood variability, are associated with negative mental health outcomes. Since adolescence is a time when mood disorder onset peaks, mood variability during this time is of significant interest. Understanding biological factors that might be associated with mood variability, such as sleep and structural brain development, could elucidate the mechanisms underlying mood and anxiety disorders. Data from the longitudinal Leiden Self-Concept study (N=171) over 5 yearly timepoints was used to study the association between sleep, brain structure, and mood variability in healthy adolescents aged 11-21 at baseline in this pre-registered study. Sleep was measured both objectively, using actigraphy, as well as subjectively, using a daily diary self-report. Negative mood variability was defined as day-to-day negative mood swings over a period of 5 days after an MRI scan. It was found that negative mood variability peaked in mid-adolescence in females while it linearly increased in males, and average negative mood showed a similar pattern. Sleep duration (subjective and objective) generally decreased throughout adolescence, with a larger decrease in males. Mood variability was not associated with sleep, but average negative mood was associated with lower self-reported energy. In addition, higher thickness in the dorsolateral prefrontal cortex (dlPFC) compared to same-age peers, suggesting a delayed thinning process, was associated with higher negative mood variability in early and mid-adolescence. Together, this study provides an insight into the development of mood variability and its association with brain structure.

## Introduction

Mood variability, or emotional instability, refers to fluctuations in mood over time^1^. Mood variability undergoes significant changes during adolescence^2^ and has been associated with negative mental health outcomes, such as an increased risk of developing anxiety and depression^3–7^. The goal of this study was to provide a comprehensive analysis of the developmental pattern of mood variability in adolescence and to examine two biological mechanisms that may account for these developmental patterns: sleep changes ^8^ and structural brain development^9^.

Mood variability has previously been studied for both positive (e.g., happiness, vigor) and negative (e.g., sadness, anger, tension) emotions^10^. Mood variability is typically assessed using daily assessment of mood states, for example through self-report questionnaires, in which participants are asked to rate their mood across several days. This allows for the assessment of both general mood as well as daily fluctuations^11^. Mood variability is defined as the average difference in mood between consecutive days. Prior studies described several developmental patterns, although different methods were used. First, one prior study described that female adolescents (10-14 years at the first time point) showed an increase in mood variability over a 4-year period, when using standard deviation to calculate mood variability, suggesting a rise in mood variability in adolescence^12^. In contrast, a second longitudinal study in adolescents starting from the age of 13 years showed that mood variability for happiness, sadness and anger linearly decreased over a five-year period in both boys and girls^13^. Furthermore, mood variability is an important outcome of emotion regulation and it was found that in mid-adolescence emotion regulation strategies were used less compared to early and late adolescence^14–16^. Thus, the results on mood variability are mixed, but it could be that mood variability peaks in mid-adolescence, possibly associated with less efficient emotion regulation strategies, followed by a decrease into adulthood.

Prior studies have suggested an important link between mood and sleep^17,18^. During the developmental period of adolescence, melatonin release shifts to a later time, leading to a later bedtime, and thereby often causing sleep deprivation^19,20^. Prior research showed that typically developing adolescents whose sleep was restricted to 6.5 hours per night reported worse emotion regulation compared to adolescents who slept almost 9 hours per night, but the relation with mood variability is still unclear^21^. Moreover, sleep problems have been identified to increase the risk for psychopathology onset later in life^22–24^. During development, total sleep duration decreases, which has been assessed using both subjective and objective measures^17,25,26^. Most prior studies examining mood and sleep were cross-sectional and included adolescents of a narrow age-range, which limits the possibility to examine developmental changes. Longitudinal studies are therefore needed to study the association between the development of ‘natural’ sleep deprivation, which adolescents often experience, and mood variability^8^. Due to the discrepancy between objective and subjective sleep measures, such as actigraphy underestimating sleep due to sleep motor activity, especially in boys due to their greater movement in their sleep, it is of importance that the association between both measures of sleep and mood variability are assessed^26^.

In addition to sleep related changes, structural brain development may be related to changes in mood and maturation of emotion regulation strategies^27^. During typical adolescent development, cortical thickness of the prefrontal cortex and other cortical regions consistently reduces^28–30^, and both delays and accelerations of this developmental process have been associated with symptoms of depression and anxiety^9,29,31,32^. Thicker cortical regions have been found in females, however, especially in early childhood^28^. Brain regions that are of particular interest for the relation to mood variability because of their involvement in emotion regulation include the prefrontal cortex (PFC), more specifically the ventrolateral (vlPFC) and dorsolateral (dlPFC) regions, as well as the anterior cingulate cortex (ACC), ventral striatum (VS), amygdala, and orbitofrontal cortex (OFC)^33–36^. Neuroscience models previously suggested that these regions do not develop simultaneously, as subcortical brain regions such as the amygdala and ventral striatum are thought to develop in a faster fashion than cortical brain regions such as the dlPFC and vlPFC^37,38^. It could be argued that this imbalance might lead to emotional instability.

Taken together, there is some evidence that mood variability is developing throughout adolescence^2,12^ and heightened levels of mood variability possibly have a negative effect on mental health^39,40^, but this question has not yet been examined using longitudinal measures across the whole range of adolescence. In addition, the association with two potential mechanisms, sleep^17^ and neural development^9^, throughout the development of mood variability remain unknown. Elucidating biological mechanisms of psychological risk-factors of mental health could contribute to our understanding of the development of mental disorders. Therefore, the present study had three aims: 1) to study the typical development of mood variability throughout adolescence in a longitudinal sample, 2) to study the relation between sleep, brain structure and mood variability throughout adolescent development, 3) to examine if mood variability during adolescence can predict anxiety and depressive symptoms.

We examined these questions in a pre-registered (https://osf.io/xmbg4/) comprehensive longitudinal study including three waves of neural development and daily variability in mood and sleep in adolescents between 11-24 years of age. We hypothesized that (a) mood variability develops in an inverted U-shape, (b) sleep decreases throughout development, (c) lower sleep length is associated with higher mood variability, (d) brain development of regions involved in emotion regulation is associated with mood variability, and (e) higher mood variability precedes symptoms of anxiety and depression.

## Results

Data from 171 participants over 5 time points were included, leading to a total of 661 observations (Table 1, Supplementary Table S1).

**Table 1.** Overview participants over 5 time points. Mean (SD) are being displayed.

|  | **Time point 1**<br>**(N=161)** | **Time point 2**<br>**(N=168)** | **Time point 3**<br>**(N=175)** | **Time point 4**<br>**(N=117)** | **Time point 5**<br>**(N=40)** |
| --- | --- | --- | --- | --- | --- |
| **Sex N (%)** |  |  |  |  |  |
| Male | 71 (44.1%) | 78 (46.4%) | 81 (46.3%) | 55 (47.0%) | 67 (47.9%) |
| Female | 83 (51.6%) | 81 (48.2%) | 88 (50.3%) | 62 (53.0%) | 73 (52.1%) |
| **Age** | 15.3 (2.92) | 16.0 (3.21) | 17.5 (3.46) | 18.9 (3.39) | 20.0 (3.72) |
| **Education N (%)** |  |  |  |  |  |
| Primary school | 30 (19.1%) | 24 (14.3%) | 7 (4.0%) |  |  |
| High school | 85 (52.8%) | 87 (51.8%) | 80 (45.7%) |  |  |
| Higher education | 39 (24.2%) | 48 (28.6%) | 83 (47.4%) |  |  |
| **Country of birth N (%)** |  |  |  |  |  |
| Netherlands | 147 (91.3%) | 152 (90.5%) | 161 (92.0%) |  |  |
| Turkey | 0 (0%) | 0 (0%) | 1 (0.6%) |  |  |
| Netherlands Antilles | 1 (0.6%) | 1 (0.6%) | 1 (0.6%) |  |  |
| Other | 6 (3.7%) | 6 (3.6%) | 7 (4.0%) |  |  |
| **Mood variability (negative)** | 8.41 (5.72) | 8.93 (6.29) | 9.39 (6.02) |  |  |
| **Average mood (negative)** | 10.8 (10.7) | 12.2 (11.3) | 13.5 (10.7) |  |  |
| **Mood variability** | 11.7 (6.26) | 12.2 (7.16) | 12.5 (6.54) |  |  |
| **Average mood** | 3.09 (12.3) | 4.08 (12.7) | 6.04 (12.7) |  |  |
| **Subjective sleep duration** | 9.26 (1.08) | 9.16 (0.94) | 9.02 (0.97) |  |  |
| **Sleep Energy** | 2.89 (0.54) | 2.83 (0.53) | 2.70 (0.58) |  |  |
| **Objective sleep duration** | 7.73 (0.86) | 7.58 (1.22) | 7.35 (1.03) |  |  |
| **Sleep Efficiency (%)** | 93.0 (5.63) | 91.8 (8.13) | 92.1 (7.61) |  |  |
| **Total anxiety** | 19.5 (11.8) | 15.6 (9.99) | 23.1 (14.4) | 26.6 (13.8) | 28.4 (17.2) |
| **Total depressive symptoms** | 5.72 (3.74) | 5.65 (3.95) | 7.10 (4.53) | 7.49 (4.65) | 9.63 (5.96) |

### Development of mood

The development of mood based on all 5 subscales of the Profiles of Mood States (POMS) has been reported in the supplementary materials. The development of mood was further examined using only the four negative subscales of the POMS (tension, depression, anger, and fatigue), as it was found that the positive subscale (vigor) was not associated with age-related changes.

First, we tested for sex differences in average mood and mood variability. This analysis showed that females had higher negative mood variability and negative average mood than males (*p*=0.02, *p*=0.02), therefore in all subsequent analyses the age by sex interaction was examined. Next, the development of negative mood variability was studied in an age by sex interaction model. The best-fit model on negative mood variability and negative average mood can be observed in Figure 2 (BIC: 2028.89, *p*<0.001, *k*=4; BIC: 2512.30, *p*=0.002).

**Figure 1.**
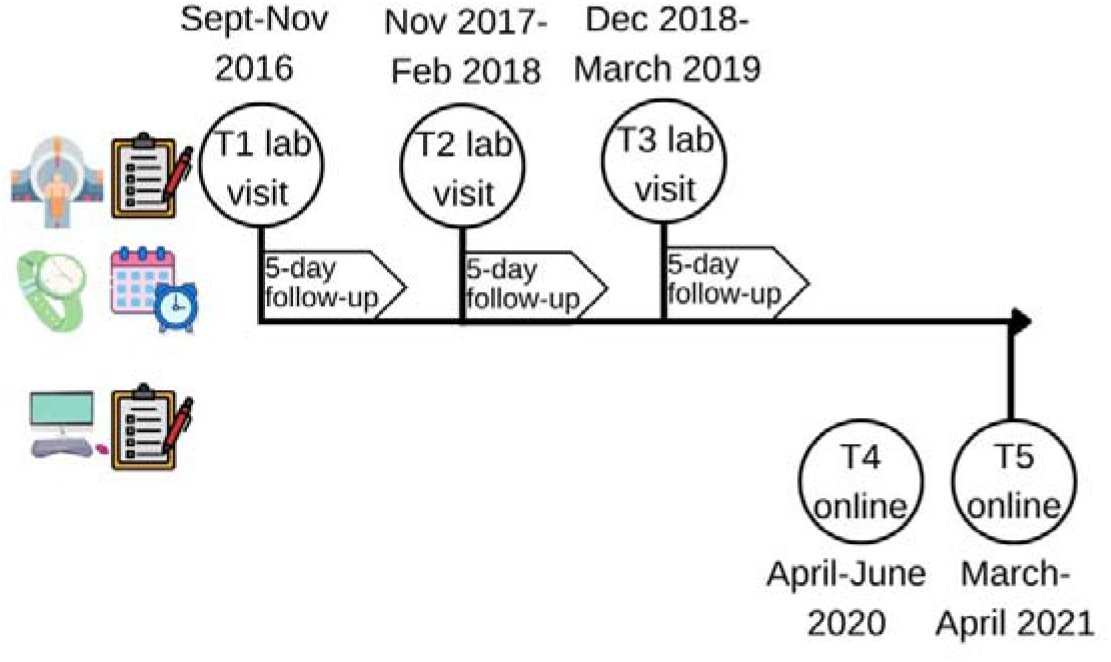
Timeline of the Leiden Self-Concept Study. At the first three time points, participants visited the lab to undergo an MRI and fill out questionnaires. In the 5 days after the lab visit, participants reported their daily mood and sleep in a diary, and sleep was measured using a wristband that detected motion. Participants were given the option to participate in two follow-up visits that only entailed filling out questionnaire measures at home.

**Figure 2.**
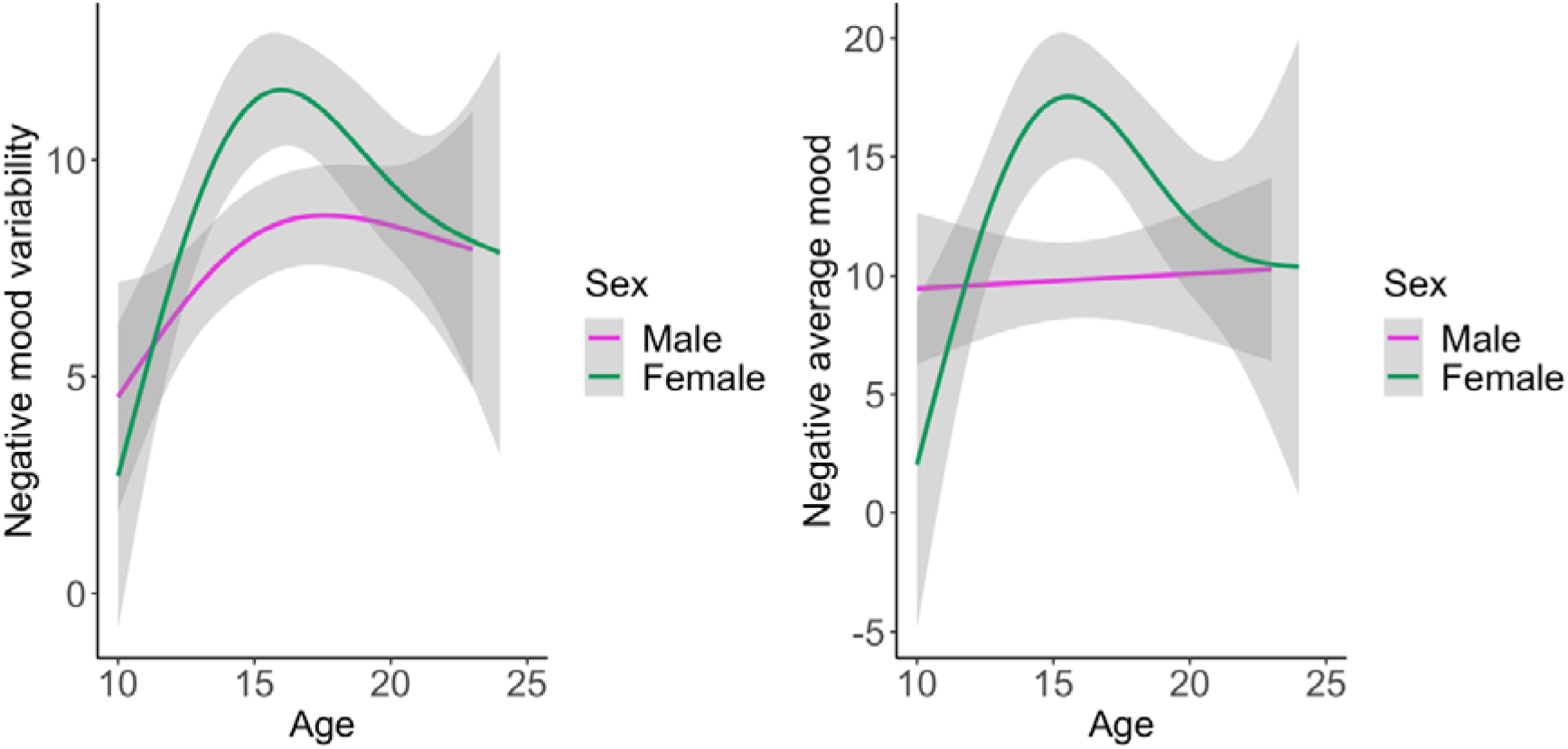
Best-fit model of negative average mood and negative mood variability development by sex. Left: negative mood variability, showing fluctuations in mood across 5 days. Right: average negative mood based in four POMS subscales (depression, anger, tension, and fatigue).

Negative mood variability showed a peak during mid-adolescence for females, showing a rapid increase and a modest decrease across adolescence (BIC: 1066.82, p=0.02, *k*=4; Figure 2 and Supplementary Figure S6). For males, negative mood variability increased throughout adolescence (BIC: 705.65, p=0.03, *k*=4). Individual trajectories of mood variability are displayed in Supplemental Figure S7. Negative average mood increased during early adolescence, also showed a peak during mid-adolescence and a decline during late adolescence for females but did not show an association with age in males (BIC: 1319.27, p<0.001, *k*=4; BIC: 1018.82, p=0.66, *k*=4).

### Development of sleep

Next, the development of sleep: objective sleep duration, objective sleep efficiency, subjective sleep duration and subjective energy level, was investigated. The best-fit model for the age by sex interaction effects are displayed in Table 2, Figure 3, and Supplementary Figure S8.

**Table 2.**
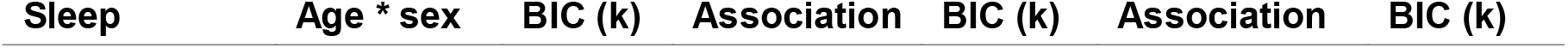
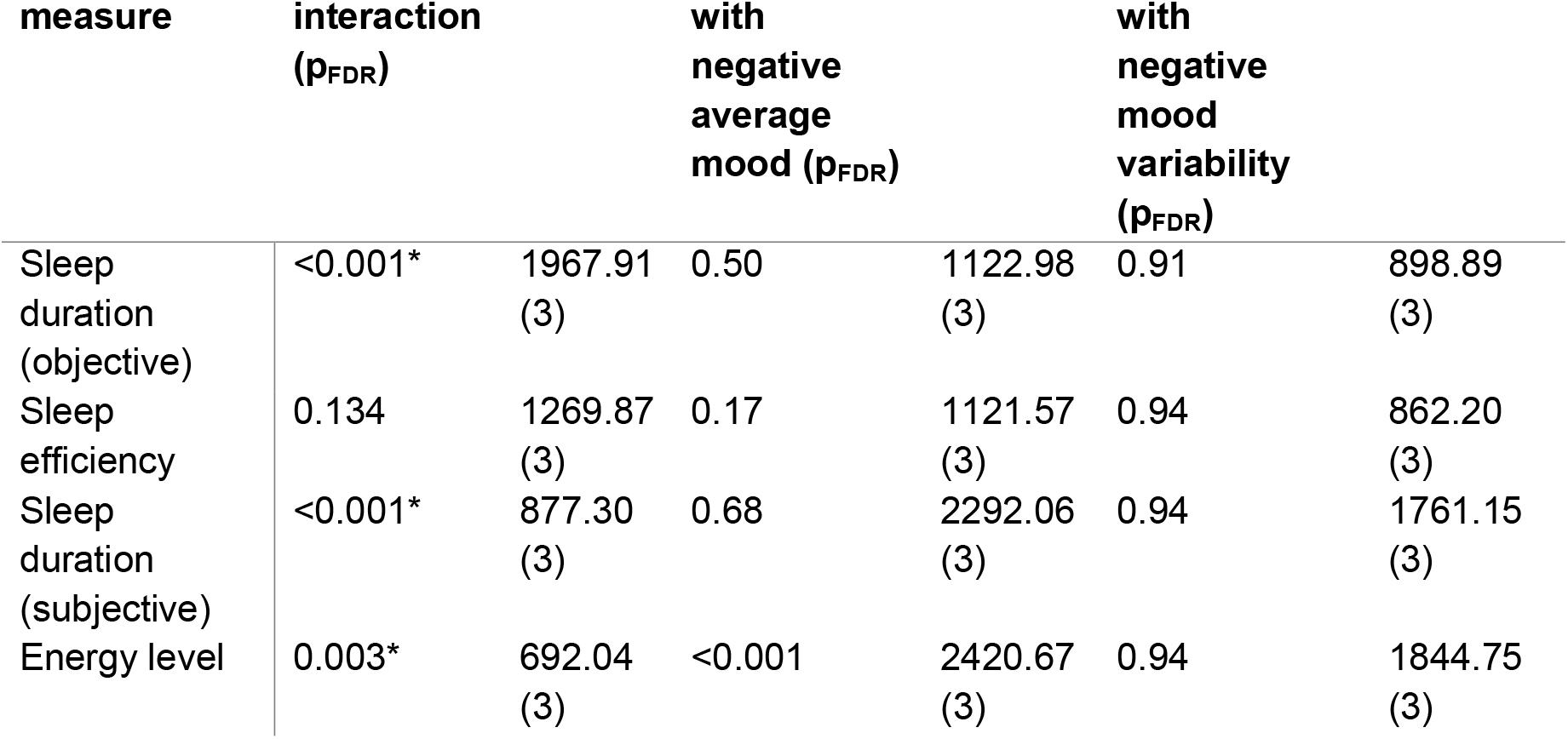
Best-fit model of sleep development.

**Figure 3.**
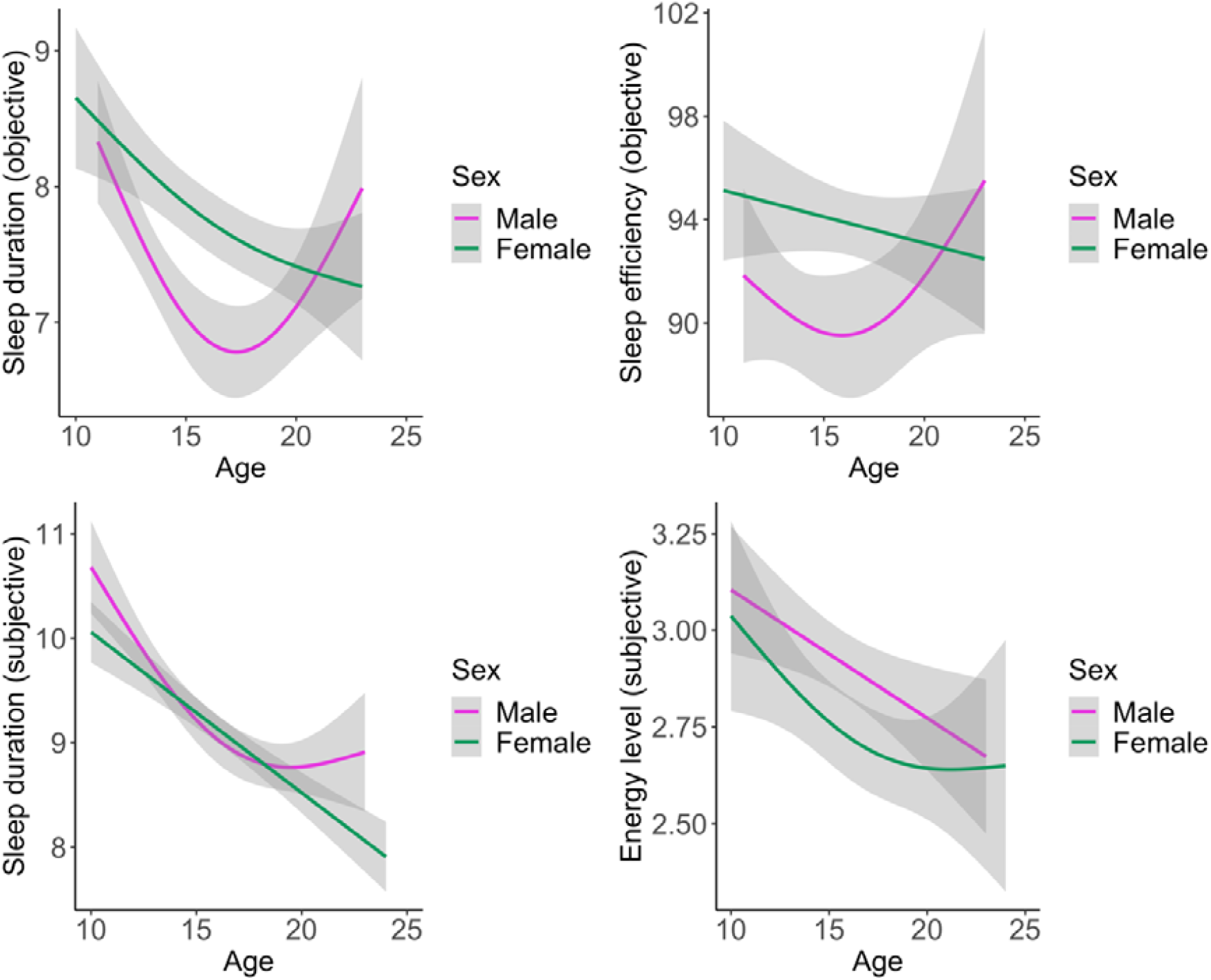
Best-fit model of sleep development by sex. Upper left: objective sleep duration, upper right: objective sleep efficiency, lower left: subjective sleep duration, lower right: energy level.

There was a significant age by sex interaction for objective and subjective sleep duration, and subjective energy level but not for sleep efficiency. Objective and subjective sleep duration decreased for both males and females, but in males there was a subsequent flattening, and even increase, in late adolescence. Objective sleep efficiency did not show an age by sex interaction. However, subjective energy level decreased throughout adolescence, for both males and females.

### Association between sleep and mood throughout adolescence

To study the association between sleep measures and mood, sleep measures were scaled to age. Sleep duration and sleep efficiency were not associated with negative mood variability or average mood. The best-fit models showed that age-corrected energy level and negative mood variability (BIC: 2483.68, p_FDR_<0.001, *k*=3) showed an association, however, this association did not survive correction for average negative mood (Table 2). A second association was also observed between age-corrected energy level and average negative mood (BIC: 2149.39, p_FDR_=0.009, *k*=3). People with lower level of energy compared to their age-matched peers showed higher negative mood throughout development (Figure 4). Subsequent control analyses confirmed that no effect of holidays was found, and the results did not change when the analyses were repeated separately for weekends and weekdays.

**Figure 4.**
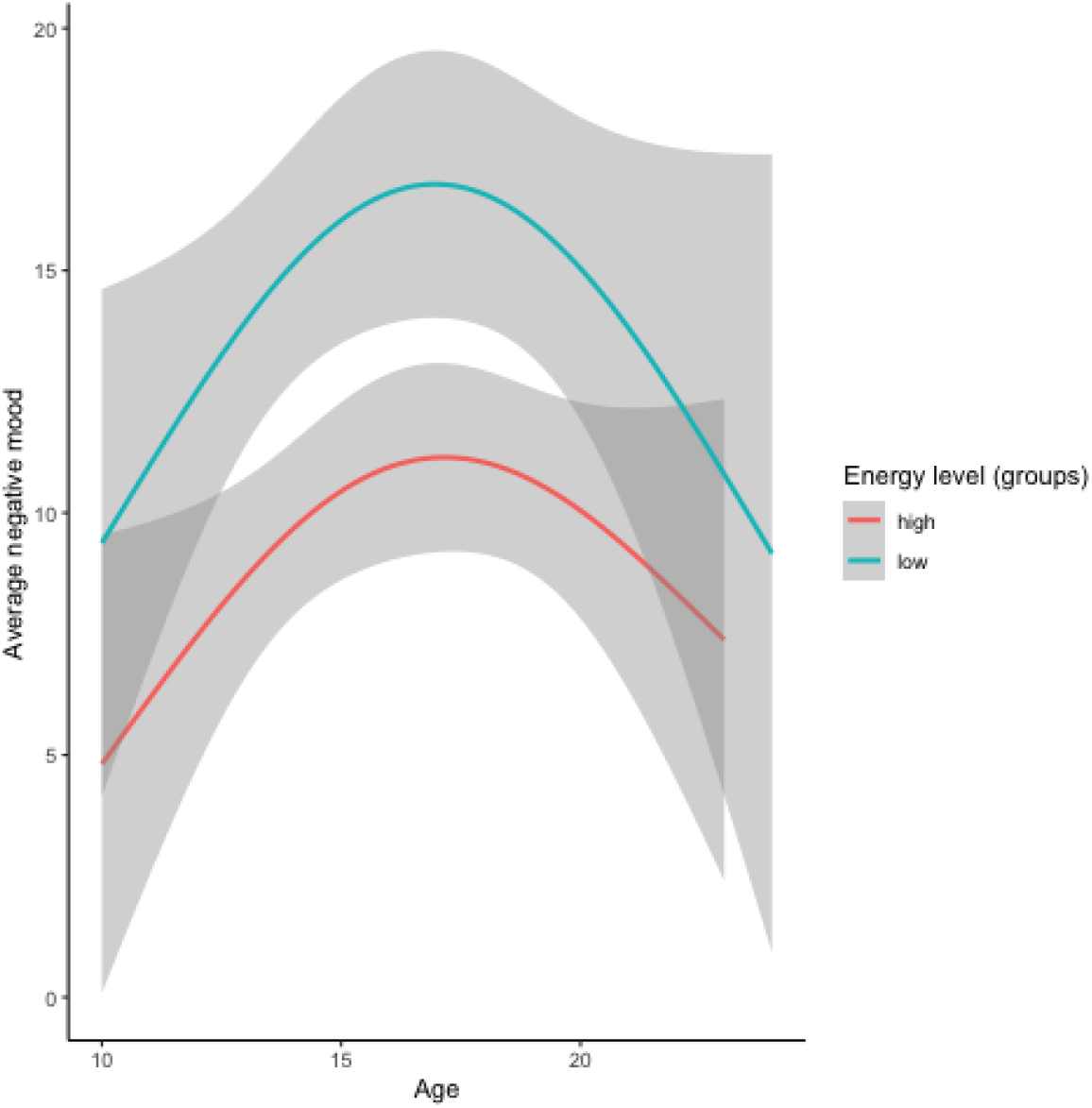
Mood variability development in groups with high or low levels of energy. (subjective measure of sleep). Participants are merely divided into groups for visualisation purposes.

### Association between brain structure and mood throughout adolescence

Next, the association between brain structure scaled to age and mood was examined. Consistent with prior studies ^41,42^, cortical thickness for all regions decreased during adolescent development (for example: dlPFC in Figure 5A). Negative mood variability showed an association with dlPFC thickness (BIC: 1792.19, p_FDR_=0.04, *k*=3). Those participants with higher dlPFC thickness compared to same-age peers showed higher levels of negative mood variability in early and mid-adolescence (Figure 5B). Negative average mood was not associated with brain structure. None of the other brain regions showed associations with negative mood variability or average mood.

**Figure 5.**
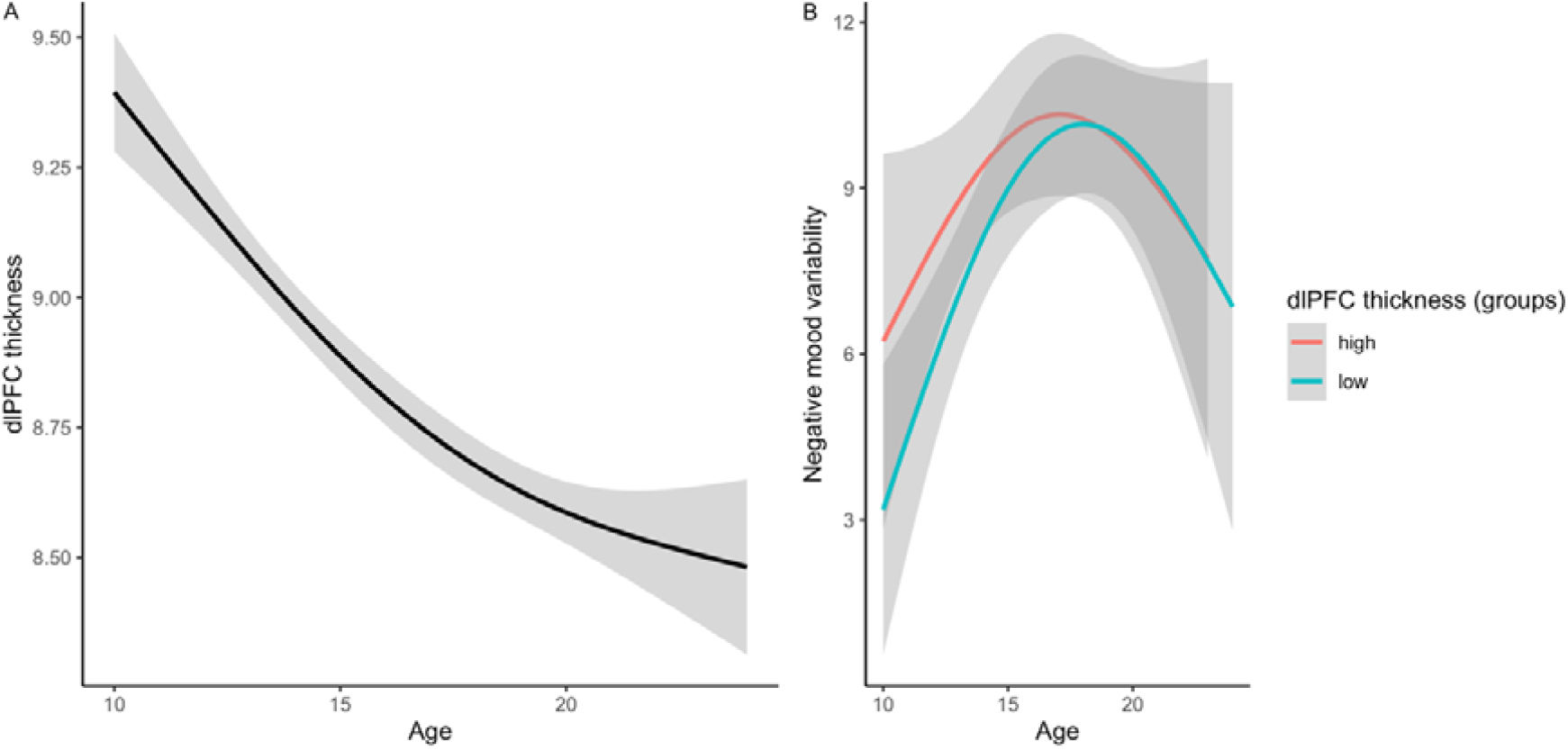
Development of dlPFC thickness (A) and mood variability development in groups with high or low dlPFC (B). Participants are merely divided into groups for visualisation purposes

### Association between mood variability and future anxiety and depressive symptomatology

The final aim was to investigate the association between mood variability and mental health, specifically whether mood variability preceded anxiety and depressive symptoms. It was found that mean mood variability at wave 1 to 3 preceded higher anxiety and depressive symptoms at wave 4 and 5 (depression: B=0.22, p=0.004; anxiety: B=1.02, p<0.001). Finally, we tested whether this effect remained after correcting for average anxiety and depressive symptoms over wave 1 to 3, but it did not survive this correction (depression: B=-0.04, p=0.61; anxiety: B=0.31, p=0.14).

## Discussion

The present study used a longitudinal design to examine the development of day-to-day mood variability during adolescent development. As predicted, negative mood variability peaked in mid-adolescence, but only for females, whereas males showed linear increases in negative mood variability with age and lower levels of mood variability overall. Objective and subjective sleep duration dropped in mid-adolescence in line with prior research ^25^ and males but not females showed a subsequent flattening and increase in sleep duration. Sleep duration was not associated with mood variability, but subjective energy levels were negatively associated with average mood (see also^17^). Lastly, consistent with prior research dorsolateral prefrontal cortex thickness declined with age^31^, and negative mood variability showed an association with the development of thickness in the dorsolateral prefrontal cortex.

Our first aim was to provide a longitudinal assessment of mood variability in adolescence. Due to the developmental patterns of mood observed (see Supplemental Material), we specifically focused on negative mood. The timing of the peak seen in mood variability was slightly later compared to previous research where a decrease was seen after age 13-years^2^. Additionally, this peak overlapped with the peak in average negative mood seen in females, showing that females not only showed larger mood swings but also an on average more negative mood in mid-adolescence. The current study adds to the literature as it completely covered adolescent development, and therefore the peak in mood variability can be observed in females. The development of mood variability was especially of interest because of its association with depression and anxiety, and onset of these psychiatric disorders peaks during adolescence^43^. However, in the current study the association between mood swings and future symptoms of depression and anxiety were explained by current mood.

The findings of a decrease in sleep duration (objective as well as subjective) fit well with prior research showing that there are sex differences in sleep development, and that overall, a decrease in sleep duration is observed^17,25^. Contrary to our predictions, neither objective nor subjective sleep duration were related to mood variability. Notably, developmental patterns associated with sex were inverted for males and females, with females showing a larger peak in mood variability and males showing a larger drop in objective sleep duration. These sex differences are currently not well understood, but some of these effects could potentially be related to social experiences and expectations. For example, males show higher levels of online gaming than females during adolescence^41^, possibly resulting in a shorter sleep duration in mid-adolescence. Alternatively, earlier research showed that adolescent females more often slept less than 6 or more than 10 hours, which could have led to a higher average than in males^25^. Lastly, previous research showed that actigraphy less accurately measures sleep in males due to movement which might contribute to the large decrease in sleep observed^26^.

Despite the absence of a relation between mood variability and sleep, lower subjective energy levels, but not objective sleep measures, were associated with lower average mood. Even though energy levels decreased during adolescence, the relationship between average mood and energy levels did not change during development, suggesting that decreasing energy levels during adolescence co-occur with generally increasing negative mood. This potentially points to a causal relationship, therefore, it could be suggested that increasing sleep duration by for example adjusting school times could be beneficial for mood. However, these questions should be studies in future research using sleep intervention designs, as the current study, despite being longitudinal, is correlational and cannot draw causal conclusions.

One alternative explanation should be considered; subjective sleep has often been more strongly associated with depressed and anxious mood than objective sleep, which might be due to low mood being associated with the misperception of sleep, meaning that when people experience low mood, they often perceive their sleep as worse^44,45^. This again highlights the need for intervention studies to explore the directionality of the association.

One of the mechanisms underlying the increased levels of mood variability in mid-adolescence might be emotion regulation^14^. Even though we did not measure emotion regulation strategies directly, we examined whether mood variability was associated with structural brain development of regions involved in emotion regulation^46^. A thicker dlPFC compared to same-age peers was associated with higher levels of negative mood variability in early and mid-adolescence, and a similar pattern was observed for mood variability and vlPFC thickness at trend level. Since adolescence is a time of cortical thinning, a thicker dlPFC could be interpreted as a delayed cortical maturation process compared to same-age peers. Thus, higher mood variability might be associated with a delay in cortical maturation. In late adolescence, after the age at which most rapid thinning of the dlPFC takes place^47^, the association with mood variability seems less pronounced. While this is the first study to show an association with mood variability, an earlier study, consistent with the current findings, showed that the development of dlPFC and vlPFC thickness was associated with the use of emotion regulation strategies, with greater cortical thinning associated with higher levels of cognitive reappraisal later in life^48^. Taken together, dlPFC thickness might be a biological marker that explains why some adolescents experience more mood swings than others.

Interestingly, no association between mood variability and limbic regions, the amygdala and ventral striatum, was found. This is however in line with work on the association between emotion regulation and subcortical development, also not reporting a direct association^49^.

One possible explanation could be that subcortical regions do not show large intra-individual brain volume differences during adolescence and therefore no relation with developmental processes of mood variability^50,51^. Differences between surface area and thickness might explain why only an association between mood variability and thickness was found. For example, thickness is more susceptible to environmental influences whereas surface area is more strongly impacted by genetic factors^52^. In future research, examining the maturational coupling or functional activity or connectivity might provide more insight into the role of the subcortical regions in mood variability^53,54^.

A few limitations of the current study should be noted. First, the subjective sleep data was subject to recall bias, and the objective data to the adolescents wearing the actigraphy watch correctly. However, the strength of assessing both subjective as well as objective reports of sleep was that the relation of these two aspects of sleep to mood variability could be compared^44^. It should be noted that we measured day-to-day variability of mood, whereas within-day variability might show a different pattern and different associations with biological mechanisms. We might not have picked up on the relation between mood variability and for example sleep because of the sampling rate of mood variability^55,56^. Future studies could examine a higher temporal resolution to measure mood variability by applying a method such as ecological momentary assessment (EMA)^57^. Additionally, future (intervention) studies should explore the causal relation between the biological mechanisms and mood variability, as from this study it remains unknown whether they precede or follow mood swing development. Finally, the relation to pubertal hormone timing and development was not assessed. Since there were sex differences in the developmental trajectory of mood variability, and mood variability increased throughout puberty in females, hormone levels might play a role. Moreover, hormones have previously been related to brain development, and pubertal timing can affect the risk for developing psychopathology in adolescence^51,58^.

In conclusion, this study confirmed the hypothesis of increased levels of mood variability in adolescence with a peak in mid-adolescence, especially in females^2,12^, and that higher levels of mood variability are associated with future symptoms of anxiety and depression^39^. We found moderate evidence for a relation between average mood and subjective sleep energy levels, and small evidence for a relation between mood variability and structural brain development. Together, these findings suggest that mood variability, sleep duration changes, and neural development are co-occurring in adolescence. Yet, the study also highlighted sex differences which may be associated with different societal expectations and experiences. Future intervention studies should explore the causal relations between the biological mechanisms identified and mood swings. This study provides initial and important insight into the normative developmental patterns and biological aspects associated with risk factors for future depressive and anxiety symptoms.

## Methods and materials

### Participants

In this study, data from adolescents from the Leiden Self-Concept Study was used^59,60^. The Leiden Self-Concept Study is a longitudinal study with 5 time points. Participants were aged 11–21 years at the first time point and recruited from schools and online. They were followed for three consecutive years with lab visits, and two additional years with online questionnaires. The lab visits were followed by 5 days in which the participants filled out questionnaires on their mood and wore an actigraphy watch. Healthy adolescents (N=171) took part in the study (Table 1, Supplementary Table S1). Participants were financially reimbursed. Exclusion criteria included the following: being left-handed, not having normal or corrected-to-normal vision, neurological or psychiatric diagnoses at T1, and usage of psychotropic medication. Participants and parents of participants younger than 18 years signed informed consent. The study (NL54510.058.16) was approved by the Medical Ethics Committee of the Leiden University Medical Centre (LUMC).

### Measures

#### Mood variability

The Profiles of Mood States (POMS) questionnaire was used to assess daily mood^61^. Participants answered these questions in an app, and they received a notification 6 pm. In this questionnaire participants were asked to rate 32 adjectives on a 5-point Likert scale to which degree the adjective described their current mood. The POMS questionnaire consists of 5 subscales: anger, depression, fatigue, tension, and vigor. The participants filled out the questionnaire daily for a five-day period after the lab visit. Mood variability was calculated per subscale as the absolute difference between successive days^13^. These scores per day were summed and then divided by the number of consecutive days the participants rated their mood. Total mood variability was defined as the sum of the mood instability on the subscales. Only participants with ratings on 3 or more consecutive days were included.

Higher scores indicate a higher mood variability. In addition, the total average mood was calculated by summing the averaged subscales and subtracting the average of the vigor subscale. Missing items were imputed using predictive mean matching from the ‘mice’ package in R^62^.

#### Sleep measures

Objective and subjective measures of sleep were included. Actigraphy watches were used to measure sleep duration and sleep efficiency objectively and using daily diaries subjective sleep duration and energy levels were assessed on the same days, for five days following the lab visit (Supplementary Methods).

#### Anxiety and depressive symptoms

During the lab visit, participants filled out the Revised Child Anxiety and Depression Scale (RCADS) to assess the participants’ feelings of anxiety and depression. Participants rated 47 questions on a Likert scale of 0–3. The answers were summed to create two subscales: total anxiety (sum of the anxiety subscales) and total depression^63^. Up to two missing items were allowed per subscale. Missing items were replaced by prorating the other items within the subscale.

#### MRI acquisition

MRI scans for the three waves were acquired on a Philips Ingenia 3.0 Tesla MR scanner. A standard whole-head coil was used. First, functional scans were obtained, followed by a high-resolution 3D T1-FFE scan (TR=9.72 ms, TE=4.6 ms, flip angle=8°, 140 slices, voxel size=0.875×0.875×0.875 mm, FOV=224×178.5×168 mm). Participants watched a film while they were in the MRI.

#### MRI processing

The MRI data was processed in the longitudinal stream in FreeSurfer 6.0 (https://surfer.nmr.mgh.harvard.edu/)^64,65^. Parcellation of the cortex was based on the Desikan-Killiany atlas and Fischl atlas for subcortical regions^66,67^. Regions of interest were selected based on prior literature on emotion regulation. Cortical thickness and surface area of the following regions were included: dlPFC, vlPFC, ACC, and OFC, as well as volume of the following regions: ventral striatum and amygdala. The average of the left and right hemisphere was used.

The regions-of-interest were constructed by combining the following regions^9,50,68,69^: dlPFC: superior frontal, rostral middle frontal cortex and caudle middle frontal, vlPFC: pars opercularis, pars triangularis and pars orbitalis, ACC: rostral ACC and caudal ACC, OFC: lateral orbitofrontal cortex and middle orbitofrontal cortex and ventral striatum: caudate, putamen and nucleus accumbens. This resulted in a total of 6 ROIs, which consisted of 4 cortical thickness, 4 cortical surface area, and 2 subcortical volume measures. The quality of the T1 images was assessed using Qoala-T^70^. Parts of these data were previously published^41,42^.

### Statistical analyses

Generalized additive mixed models (GAMMS) were used to study the development of mood variability and its association with sleep and brain structure. Generalized additive mixed models (GAMMs) are semi-parametric models that use penalized smoothing splines^71^ (Supplementary Methods).

First, an exploratory analysis was performed on the subscales (tension, anger, depression, fatigue, and vigor) to test for developmental patterns. Based on these findings, we created separate models for negative mood variability and average negative mood score, based on the first four subscales (see Supplementary Material). The main analyses (analyses 1 to 4 described below) were therefore done using these negative mood measures, as well as the general mood variability and average mood.

Data from the first three timepoints were used in the first four analyses, to study:

1. the development of mood variability (by testing the effect of age by sex on mood variability).

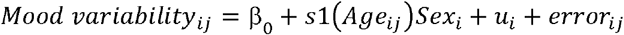
2. the development of sleep (by testing the effect of age by sex on sleep (separately for sleep duration, sleep efficiency, subjective sleep duration, energy level)).

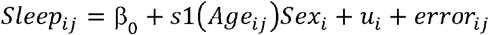
3. the association between mood variability and sleep (separately for each sleep measure, scaled by age) over time corrected for age and sex:

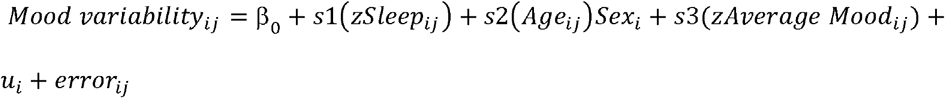
4. the association between mood variability and brain structure (scaled by age) over time corrected for age and sex.

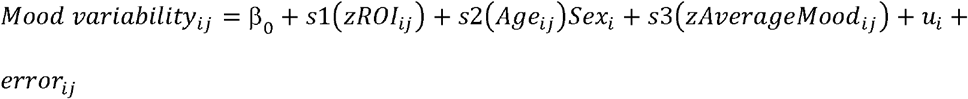

The same analyses were repeated for average mood. In each analysis, i = subject, j = time point and ui = random effect per subject. The analyses for mood variability and average mood were FDR corrected per analysis (1 to 4 above) to correct for multiple testing (so for example the association between mood variability and sleep was corrected for four tests). For analysis 4 it was corrected per type of brain measure (volume, surface area and thickness).

A thin plate spline was used to fit the data. Model fit, to choose the best *k*, was assessed using the BIC. *k* represents the basis dimensions, the maximum degrees of freedom for a smoothing spline. The ‘mgcv’ package in R was used for the analyses^72^. Participants were added as a random effect and restricted maximum likelihood (REML) was used for smoothness selection. The interaction effect of age and sex was included to test whether the development of mood variability differed between sexes. In analyses 3 and 4, sleep and brain structure were transformed into z-scores by scaling them to age. Participants with a large ROI or longer sleep duration at a certain age compared to their same-age peers had a higher z-score.

Several control analyses were conducted. The analyses for mood variability were corrected for average mood^2,11^. In addition, for analyses 2 and 3, the analyses were repeated with an additional covariate to correct for whether the participant was on holiday or at school/work.

The analyses with objective sleep measurements were also repeated separately for weekend and weekdays.

To study the prediction of anxiety and depressive symptoms based on the development of mood variability, a linear regression was used to predict average symptoms at follow-up (averaged over wave 4 and 5) based on the average level of mood variability^73,74^. The analysis was corrected for the baseline level of anxiety and depressive symptoms by adding the mean level of anxiety or depressive symptoms at the first three waves as a covariate.

## Acknowledgments

This work was supported by the Netherlands Organisation for Scientific Research (NWO-VICI 453-14-001 E.A.C.). We thank all participants to the Leiden Self-Concept study.

## Data Availability

The data used is the study is available upon request.

## Author Contributions

Y.T. and E.C. conceptualized the study. R.C. and S.G. collected the data. Y.T. performed the analyses, with contribution from L.W. and E.C. Y.T. wrote the main manuscript with contribution from E.C. All authors reviewed and approved the final manuscript.

## References

1. Bailen, N. H., Green, L. M. & Thompson, R. J. Understanding Emotion in Adolescents: A Review of Emotional Frequency, Intensity, Instability, and Clarity. Emotion Review 11, 63–73 (2019).

2. Maciejewski, D. F., van Lier, P. A. C., Branje, S. J. T., Meeus, W. H. J. & Koot, H. M. A 5-Year Longitudinal Study on Mood Variability Across Adolescence Using Daily Diaries. Child Dev 86, 1908–1921 (2015).

3. Maciejewski, D. F. et al. The development of adolescent generalized anxiety and depressive symptoms in the context of adolescent mood variability and parent-adolescent negative interactions. J Abnorm Child Psychol 42, 515–526 (2014).

4. Neumann, A., Van Lier, P. A. C., Frijns, T., Meeus, W. & Koot, H. M. Emotional dynamics in the development of early adolescent psychopathology: A one-year longitudinal study. J Abnorm Child Psychol 39, 657–669 (2011).

5. Silk, J. S., Steinberg, L. & Morris, A. S. Adolescents’ Emotion Regulation in Daily Life: Links to Depressive Symptoms and Problem Behavior. Child Dev 74, 1869–1880 (2003).

6. Koval, P., Sütterlin, S. & Kuppens, P. Emotional inertia is associated with lower well-being when controlling for differences in emotional context. Front Psychol 6, (2016).

7. Kuppens, P. et al. Emotional inertia prospectively predicts the onset of depressive disorder in adolescence. Emotion 12, 283–289 (2012).

8. Galván, A. The Need for Sleep in the Adolescent Brain. Trends Cogn Sci 24, 79–89 (2020).

9. Bos, M. G. N. et al. Longitudinal structural brain development and externalizing behavior in adolescence. J Child Psychol Psychiatry 59, 1061–1072 (2018).

10. Green, K. H. et al. Mood and emotional reactivity of adolescents during the COVID-19 pandemic: short-term and long-term effects and the impact of social and socioeconomic stressors. Sci Rep 11, (2021).

11. Ebner-Priemer, U. W., Eid, M., Kleindienst, N., Stabenow, S. & Trull, T. J. Analytic Strategies for Understanding Affective (In)Stability and Other Dynamic Processes in Psychopathology. J Abnorm Psychol 118, 195–202 (2009).

12. Larson, R. W., Moneta, G., Richards, M. H. & Wilson, S. Continuity, Stability, and Change in Daily Emotional Experience across Adolescence. Child Dev 73, 1151–1165 (2002).

13. Maciejewski, D. F., van Lier, P. A. C., Branje, S. J. T., Meeus, W. H. J. & Koot, H. M. A 5-Year Longitudinal Study on Mood Variability Across Adolescence Using Daily Diaries. Child Dev 86, 1908–1921 (2015).

14. Zimmermann, P. & Iwanski, A. Emotion regulation from early adolescence to emerging adulthood and middle adulthood: Age differences, gender differences, and emotion-specific developmental variations. Int J Behav Dev 38, 182–194 (2014).

15. Hoeksma, J. B., Oosterlaan, J. & Schipper, E. M. Emotion Regulation and the Dynamics of Feelings: A Conceptual and Methodological Framework.

16. Cole, P. M. & Hall, S. E. Emotion dysregulation as a risk factor for psychopathology. Child and Adolescent Psychopathology (2008).

17. Short, M. A., Booth, S. A., Omar, O., Ostlundh, L. & Arora, T. The relationship between sleep duration and mood in adolescents: A systematic review and meta-analysis. Sleep Medicine Reviews vol. 52 Preprint at https://doi.org/10.1016/j.smrv.2020.101311 (2020).

18. Gillett, G., Watson, G., Saunders, K. E. & McGowan, N. M. Sleep and circadian rhythm actigraphy measures, mood instability and impulsivity: A systematic review. Journal of Psychiatric Research vol. 144 66–79 Preprint at https://doi.org/10.1016/j.jpsychires.2021.09.043 (2021).

19. Galván, A. The Need for Sleep in the Adolescent Brain. Trends Cogn Sci 24, 79–89 (2020).

20. Hagenauer, M. H., Perryman, J. I., Lee, T. M. & Carskadon, M. A. Adolescent changes in the homeostatic and circadian regulation of sleep. Dev Neurosci 31, 276–284 (2009).

21. Baum, K. T. et al. Sleep restriction worsens mood and emotion regulation in adolescents. J Child Psychol Psychiatry 55, 180–190 (2014).

22. Ong, S. H., Wickramaratne, P., Min Tang & Weissman, M. M. Early childhood sleep and eating problems as predictors of adolescent and adult mood and anxiety disorders. J Affect Disord 96, 1–8 (2006).

23. Gregory, A. M. et al. Prospective longitudinal associations between persistent sleep problems in childhood and anxiety and depression disorders in adulthood. J Abnorm Child Psychol 33, 157–163 (2005).

24. Hertenstein, E. et al. Insomnia as a predictor of mental disorders: A systematic review and meta-analysis. Sleep Medicine Reviews vol. 43 96–105 Preprint at https://doi.org/10.1016/j.smrv.2018.10.006 (2019).

25. Maslowsky, J. & Ozer, E. J. Developmental trends in sleep duration in adolescence and young adulthood: Evidence from a national United States sample. Journal of Adolescent Health 54, 691–697 (2014).

26. Short, M. A., Gradisar, M., Lack, L. C., Wright, H. & Carskadon, M. A. The discrepancy between actigraphic and sleep diary measures of sleep in adolescents. Sleep Med 13, 378–384 (2012).

27. Silvers, J. A. Adolescence as a pivotal period for emotion regulation development for consideration at current opinion in psychology. Current Opinion in Psychology vol. 44 258–263 Preprint at https://doi.org/10.1016/j.copsyc.2021.09.023 (2022).

28. Wierenga, L. M., Langen, M., Oranje, B. & Durston, S. Unique developmental trajectories of cortical thickness and surface area. Neuroimage 87, 120–126 (2014).

29. Shaw, P. et al. Neurodevelopmental trajectories of the human cerebral cortex. Journal of Neuroscience 28, 3586–3594 (2008).

30. Tamnes, C. K. et al. Development of the Cerebral Cortex across Adolescence: A Multisample Study of Inter-Related Longitudinal Changes in Cortical Volume, Surface Area, and Thickness. The Journal of Neuroscience 37, 3402–3412 (2017).

31. Tamnes, C. K. et al. Brain maturation in adolescence and young adulthood: Regional age-related changes in cortical thickness and white matter volume and microstructure. Cerebral Cortex 20, 534–548 (2010).

32. Bos, M. G. N., Peters, S., van de Kamp, F. C., Crone, E. A. & Tamnes, C. K. Emerging depression in adolescence coincides with accelerated frontal cortical thinning. J Child Psychol Psychiatry 59, 994–1002 (2018).

33. Ahmed, S. P., Bittencourt-Hewitt, A. & Sebastian, C. L. Neurocognitive bases of emotion regulation development in adolescence. Dev Cogn Neurosci 15, 11–25 (2015).

34. Blakemore, S. J. The social brain in adolescence. Nature Reviews Neuroscience vol. 9 267–277 Preprint at https://doi.org/10.1038/nrn2353 (2008).

35. Silvers, J. A., Buhle, J. T., Ochsner, K. N. & Silvers, J. The neuroscience of emotion regulation: Basic mechanisms and their role in development, aging and psychopathology. in The handbook of cognitive neuroscience (Oxford University Press, 2014).

36. Martin, R. E. & Ochsner, K. N. The neuroscience of emotion regulation development: Implications for education. Current Opinion in Behavioral Sciences vol. 10 142–148 Preprint at https://doi.org/10.1016/j.cobeha.2016.06.006 (2016).

37. Casey, B. J., Getz, S. & Galvan, A. The adolescent brain. Developmental Review 28, 62–77 (2008).

38. Somerville, L. H. & Casey, B. J. Developmental neurobiology of cognitive control and motivational systems. Current Opinion in Neurobiology vol. 20 236–241 Preprint at https://doi.org/10.1016/j.conb.2010.01.006 (2010).

39. Maciejewski, D. F. et al. The development of adolescent generalized anxiety and depressive symptoms in the context of adolescent mood variability and parent-adolescent negative interactions. J Abnorm Child Psychol 42, 515–526 (2014).

40. Neumann, A., van Lier, P. A. C., Frijns, T., Meeus, W. & Koot, H. M. Emotional dynamics in the development of early adolescent psychopathology: A one-year longitudinal study. J Abnorm Child Psychol 39, 657–669 (2011).

41. Achterberg, M. et al. Longitudinal associations between social media use, mental well-being and structural brain development across adolescence. Dev Cogn Neurosci 54, (2022).

42. Becht, A. I. et al. Longitudinal associations between structural prefrontal cortex and nucleus accumbens development and daily identity formation processes across adolescence. Dev Cogn Neurosci 46, (2020).

43. Kessler, R. C. et al. Lifetime Prevalence and Age-of-Onset Distributions of DSM-IV Disorders in the National Comorbidity Survey Replication. Arch Gen Psychiatry 62, 593–602 (2005).

44. Difrancesco, S. et al. Sleep, circadian rhythm, and physical activity patterns in depressive and anxiety disorders: A 2-week ambulatory assessment study. Depress Anxiety 36, 975–986 (2019).

45. Herbert, V., Pratt, D., Emsley, R. & Kyle, S. D. Predictors of nightly subjective-objective sleep discrepancy in poor sleepers over a seven-day period. Brain Sci 7, (2017).

46. Etkin, A., Büchel, C. & Gross, J. J. The neural bases of emotion regulation. Nature Reviews Neuroscience vol. 16 693–700 Preprint at https://doi.org/10.1038/nrn4044 (2015).

47. Fuhrmann, D., Madsen, K. S., Johansen, L. B., Baaré, W. F. C. & Kievit, R. A. The midpoint of cortical thinning between late childhood and early adulthood differs between individuals and brain regions: Evidence from longitudinal modelling in a 12-wave neuroimaging sample. Neuroimage 261, (2022).

48. Vijayakumar, N. et al. Thinning of the lateral prefrontal cortex during adolescence predicts emotion regulation in females. Soc Cogn Affect Neurosci 9, 1845–1854 (2014).

49. Ferschmann, L. et al. Cognitive reappraisal and expressive suppression relate differentially to longitudinal structural brain development across adolescence. Cortex 136, 109–123 (2021).

50. Mills, K. L., Goddings, A. L., Clasen, L. S., Giedd, J. N. & Blakemore, S. J. The developmental mismatch in structural brain maturation during adolescence. Dev Neurosci 36, 147–160 (2014).

51. Wierenga, L. M. et al. Unraveling age, puberty and testosterone effects on subcortical brain development across adolescence. Psychoneuroendocrinology 91, 105–114 (2018).

52. Fjell, A. M. et al. Continuity and Discontinuity in Human Cortical Development and Change from Embryonic Stages to Old Age. Cerebral Cortex 29, 3879–3890 (2019).

53. Vijayakumar, N. et al. Cortico-amygdalar maturational coupling is associated with depressive symptom trajectories during adolescence. Neuroimage 156, 403–411 (2017).

54. Gee, D. G. et al. A developmental shift from positive to negative connectivity in human amygdala-prefrontal circuitry. Journal of Neuroscience 33, 4584–4593 (2013).

55. Tavernier, R., Choo, S. B., Grant, K. & Adam, E. K. Daily affective experiences predict objective sleep outcomes among adolescents. J Sleep Res 25, 62–69 (2016).

56. Parsons, C. E., Schofield, B., Batziou, S. E., Ward, C. & Young, K. S. Sleep quality is associated with emotion experience and adaptive regulation of positive emotion: An experience sampling study. J Sleep Res (2021) doi:10.1111/jsr.13533.

57. Baltasar-Tello, I., Miguélez-Fernández, C., Peñuelas-Calvo, I. & Carballo, J. J. Ecological Momentary Assessment and Mood Disorders in Children and Adolescents: a Systematic Review. Current Psychiatry Reports vol. 20 Preprint at https://doi.org/10.1007/s11920-018-0913-z (2018).

58. Sisk, C. L. & Zehr, J. L. Pubertal hormones organize the adolescent brain and behavior. Frontiers in Neuroendocrinology vol. 26 163–174 Preprint at https://doi.org/10.1016/j.yfrne.2005.10.003 (2005).

59. van der Cruijsen, R., Peters, S., van der Aar, L. P. E. & Crone, E. A. The neural signature of self-concept development in adolescence: The role of domain and valence distinctions. Dev Cogn Neurosci 30, 1–12 (2018).

60. Crone, E., Green, K., van de Groep, I. & van der Cruijsen, R. A neurocognitive model of self-concept development in adolescence. under review (2022).

61. Curran, S. L., Andrykowski, M. A. & Studts, J. L. Short form of the Profile of Mood States (POMS-SF): Psychometric information. Psychol Assess 7, 80–83 (1995).

62. van Buuren, S. & Groothuis-Oudshoorn, K. Journal of Statistical Software mice: Multivariate Imputation by Chained Equations in R. vol. 45 http://www.jstatsoft.org/ (2011).

63. Chorpita, B. F., Yim, L., Moffitt, C., Umemoto, L. A. & Francis, S. E. Assessment of symptoms of DSM-IV anxiety and depression in children: a revised child anxiety and depression scale. Behaviour Research and Therapy 38, 835–855 (2000).

64. Dale, A. M., Fischl, B. & Sereno, M. I. Cortical surface-based analysis. I. Segmentation and surface reconstruction. Neuroimage 9, 179–194 (1999).

65. Reuter, M., Schmansky, N. J., Rosas, H. D. & Fischl, B. Within-subject template estimation for unbiased longitudinal image analysis. Neuroimage 61, 1402–1418 (2012).

66. Desikan, R. S. et al. An automated labeling system for subdividing the human cerebral cortex on MRI scans into gyral based regions of interest. Neuroimage 31, 968–980 (2006).

67. Fischl, B. et al. Whole brain segmentation: Automated labeling of neuroanatomical structures in the human brain. Neuron 33, 341–355 (2002).

68. Quan, M. et al. White matter tract abnormalities between rostral middle frontal gyrus, inferior frontal gyrus and striatum in first-episode schizophrenia. Schizophr Res 145, 1–10 (2013).

69. Bentley, J. N. et al. Subcortical Intermittent Theta-Burst Stimulation (iTBS) Increases Theta-Power in Dorsolateral Prefrontal Cortex (DLPFC). Front Neurosci 14, (2020).

70. Klapwijk, E. T., van de Kamp, F., van der Meulen, M., Peters, S. & Wierenga, L. M. Qoala-T: A supervised-learning tool for quality control of FreeSurfer segmented MRI data. Neuroimage 189, 116–129 (2019).

71. Wood, S. N. Stable and efficient multiple smoothing parameter estimation for generalized additive models. J Am Stat Assoc 99, 673–686 (2004).

72. Wood, S. N. Generalized additive models: An introduction with R, second edition. Generalized Additive Models: An Introduction with R, Second Edition (2017). doi:10.1201/9781315370279.

73. Wierenga, L. M., Bos, M. G. N., van Rossenberg, F. & Crone, E. A. Sex effects on development of brain structure and executive functions: Greater variance than mean effects. J Cogn Neurosci 31, 730–753 (2019).

74. Khundrakpam, B. S. et al. Exploring Individual Brain Variability during Development based on Patterns of Maturational Coupling of Cortical Thickness: A Longitudinal MRI Study. Cerebral Cortex 29, 178–188 (2019).

