## Supplementary Material for "Mood variability during adolescent development and its relation to sleep and brain development" (from another version of this preprint: version 1, 2022-08-25)

### Supplementary Methods

#### *Objective sleep*

Objective sleep was measured using actigraphy. A wristband was worn by the participants on the five consecutive days after the annual lab visit. The wristband detects gross motor activity, and this information can be used to monitor sleep. The motion data was collected in 60 seconds time frames using the MicroMini-Motionlogger actigraphy from Ambulatory Monitoring Inc. (Ardsley, NY) in zero-crossing mode. Using the zero-crossing mode, all activity above the threshold (0) within one epoch (60 seconds) was counted. The average sleep duration was calculated over the five days after the lab visit. In addition, average sleep efficiency over the five days was calculated. Sleep efficiency was defined as the percentage of the total sleep time while being in bed. The actigraphy data was analysed using the Action-W software (Version 3.68.0.1). The data was manually checked following the Action-W user guidelines and the Sadeh algorithm was used to compute the sleep duration and sleep efficiency (1). Nights with less than four hours of recorded sleep were excluded (2). Participants with actigraphy data on four or more days were included (3,4). In an exploratory analysis, the actigraphy data from weekdays and weekends was analysed separately since school rhythm can affect sleep duration in adolescents (5).

#### *Subjective sleep*

Subjective sleep was measured using the participants' daily diary. Total time in bed (time to bed till time of getting up) and energy level during the day (as a measure of sleep quality) were being included. Energy level during the day was rated on a 5-point scale (from 1 – 'Little energy or motivation to do something' to 5 – 'Unusually energised and hyperactive') (6). Energy level was used as a proxy measure for sleep quality.

#### *Statistical analysis*

GAMMs can capture non-linear growth during development and the splines are penalised to prevent overfitting. Because of the longitudinal accelerated design, meaning that participants differed in their age at the start of the study, a statistical model that can take the within-subject design into account was needed (60). In addition, mood variability, brain structure and sleep are developing throughout adolescence, and therefore the model needed to capture this potential non-linear development.

### Supplementary Results

The development of negative mood variability was assessed over 341 data points from 164 unique participants over 3 time points (Supplementary Table S1). The development of negative average mood and negative mood variability including individual patterns are displayed in Supplemental Figure S1.

**Supplementary Table S1. Overview of data available per measure**

| Measure | Observations | Unique participants |
|---|---|---|
| Mood variability | 341 | 164 |
| Sleep duration (objective) | 173 | 105 |
| Sleep efficiency | 173 | 105 |
| Sleep duration (subjective) | 270 | 146 |
| Energy level | 290 | 149 |
| Brain structure | 334 | 162 |

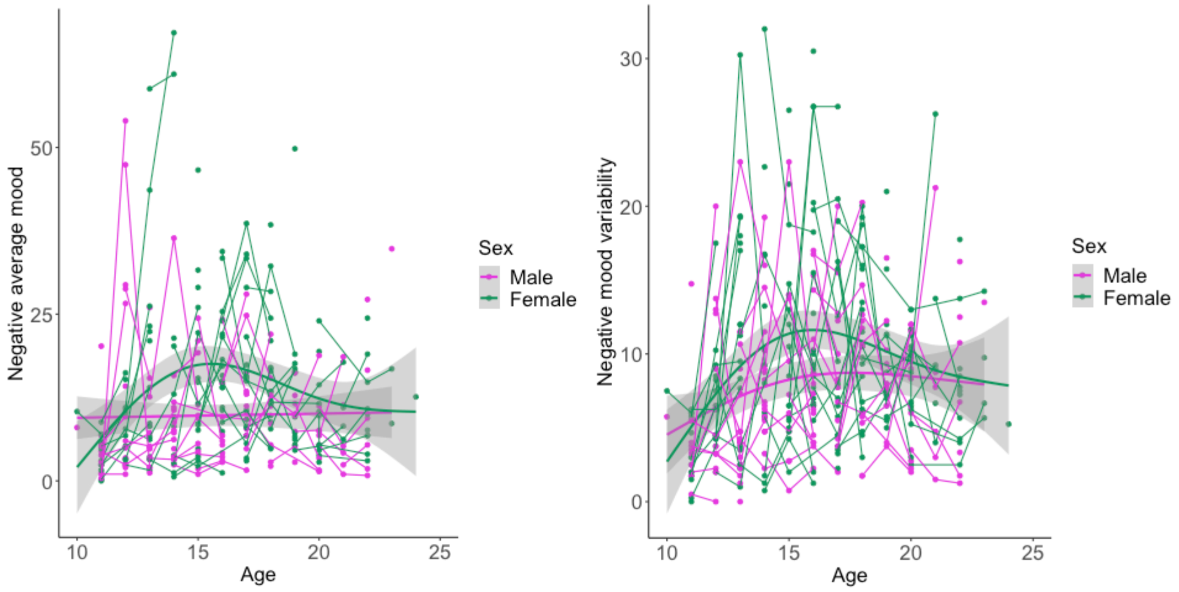


**Supplementary Figure S1. Best-fit model of negative average mood and negative mood variability development by sex.** Left: average negative mood, Right: negative mood variability.

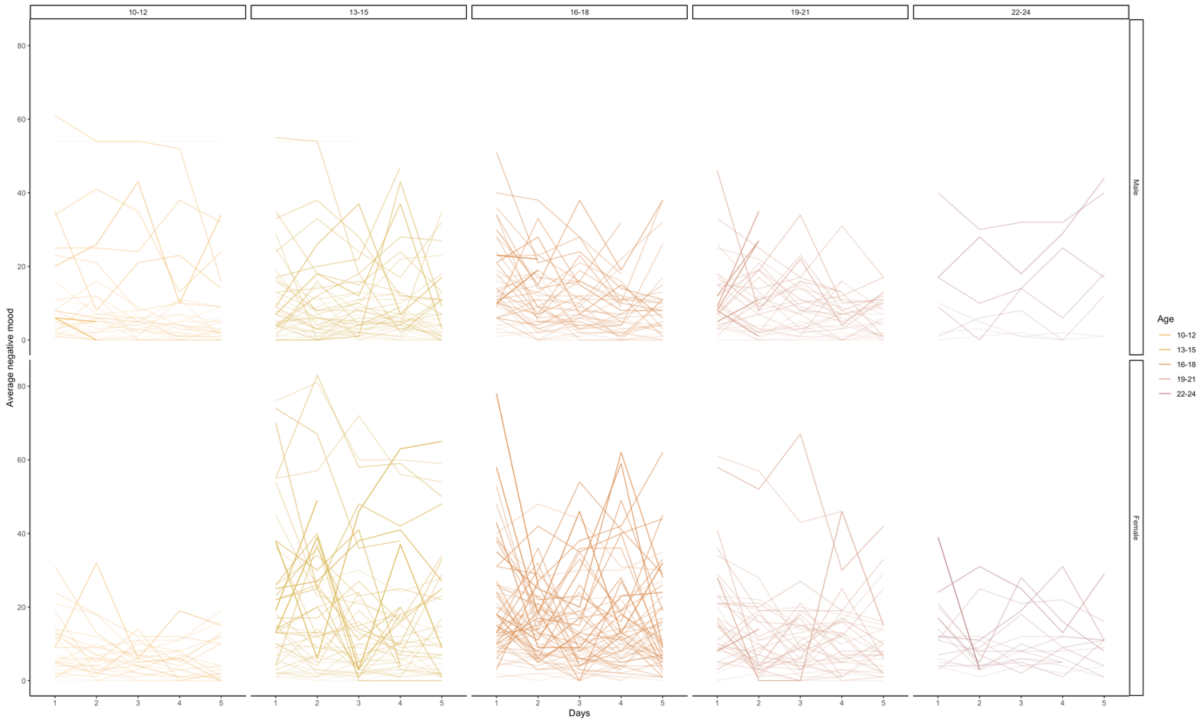


**Supplementary Figure S2. Individual trajectories of negative mood.** The variability of the lines reflect the variability of negative mood. The lines are coloured by their level of mood variability and age. Upper panel: females, lower panel: males.

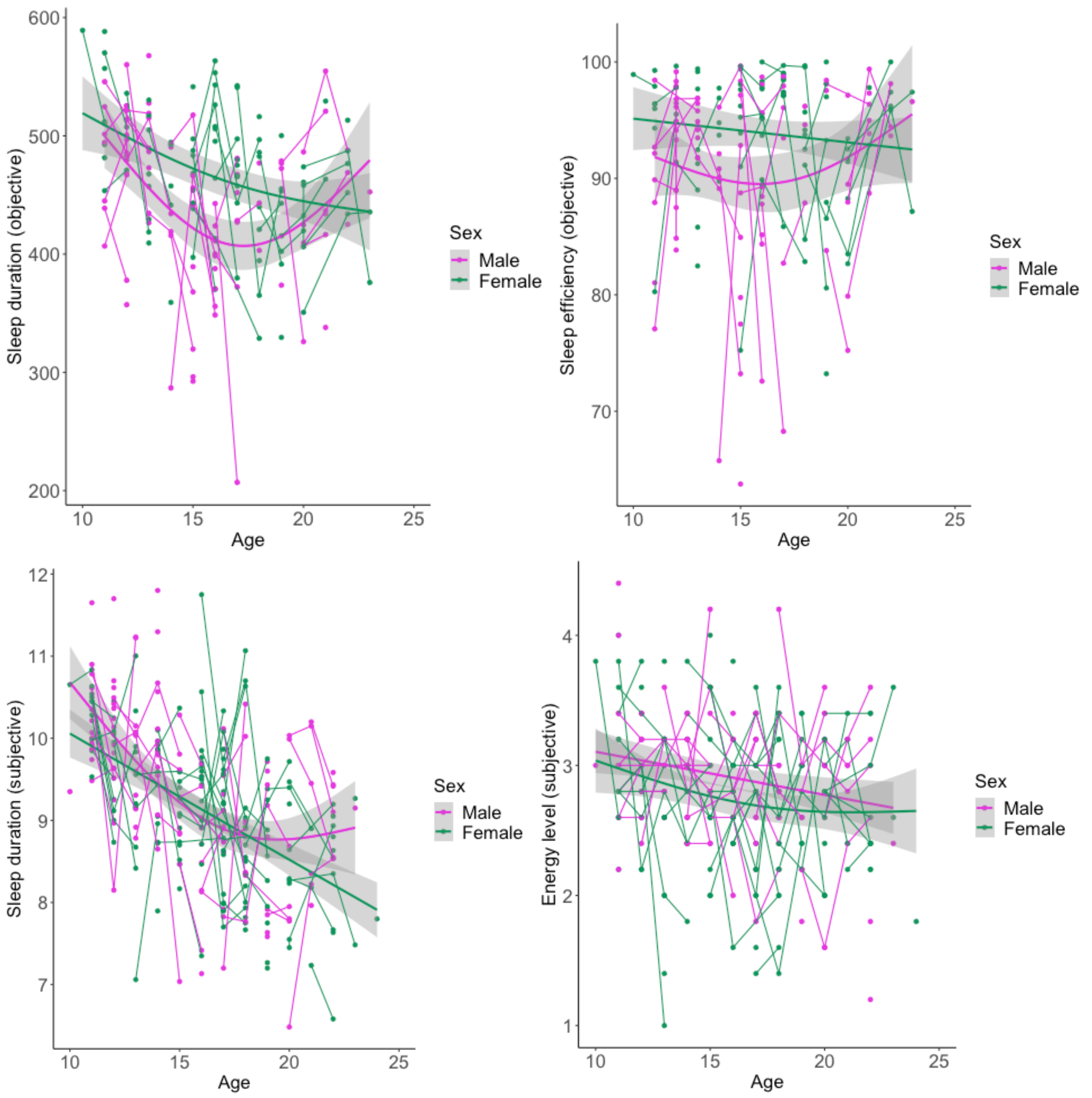


**Supplementary Figure S3. Best-fit model of sleep development by sex**. Upper left: objective sleep duration, upper right: objective sleep efficiency, lower left: subjective sleep duration, lower right: energy level.

*Development of mood per subscale*

The development of mood variability was assessed exploratively separately per subscale as earlier research showed different developmental trajectories per subscale. Variability on the four negative mood subscales, tension, anger, fatigue, and depression, but not the positive mood subscale, vigor, showed an interaction effect between age and sex (Supplementary Table S2, Supplementary Figure S4). The same was found for average mood, with anger trending towards significance (Supplementary Table S2, Supplementary Figure S5).

**Supplementary Table S2. Development of mood per subscale.**

| **Subscale** | **Mood variability $p_{FDR}$** | **Average mood $p_{FDR}$** |
|---|---|---|
| Tension | <0.001 | 0.047 |
| Depression | <0.001 | 0.047 |
| Anger | <0.001 | 0.063 |
| Fatigue | <0.001 | 0.004 |
| Vigor | 0.21 | 0.17 |

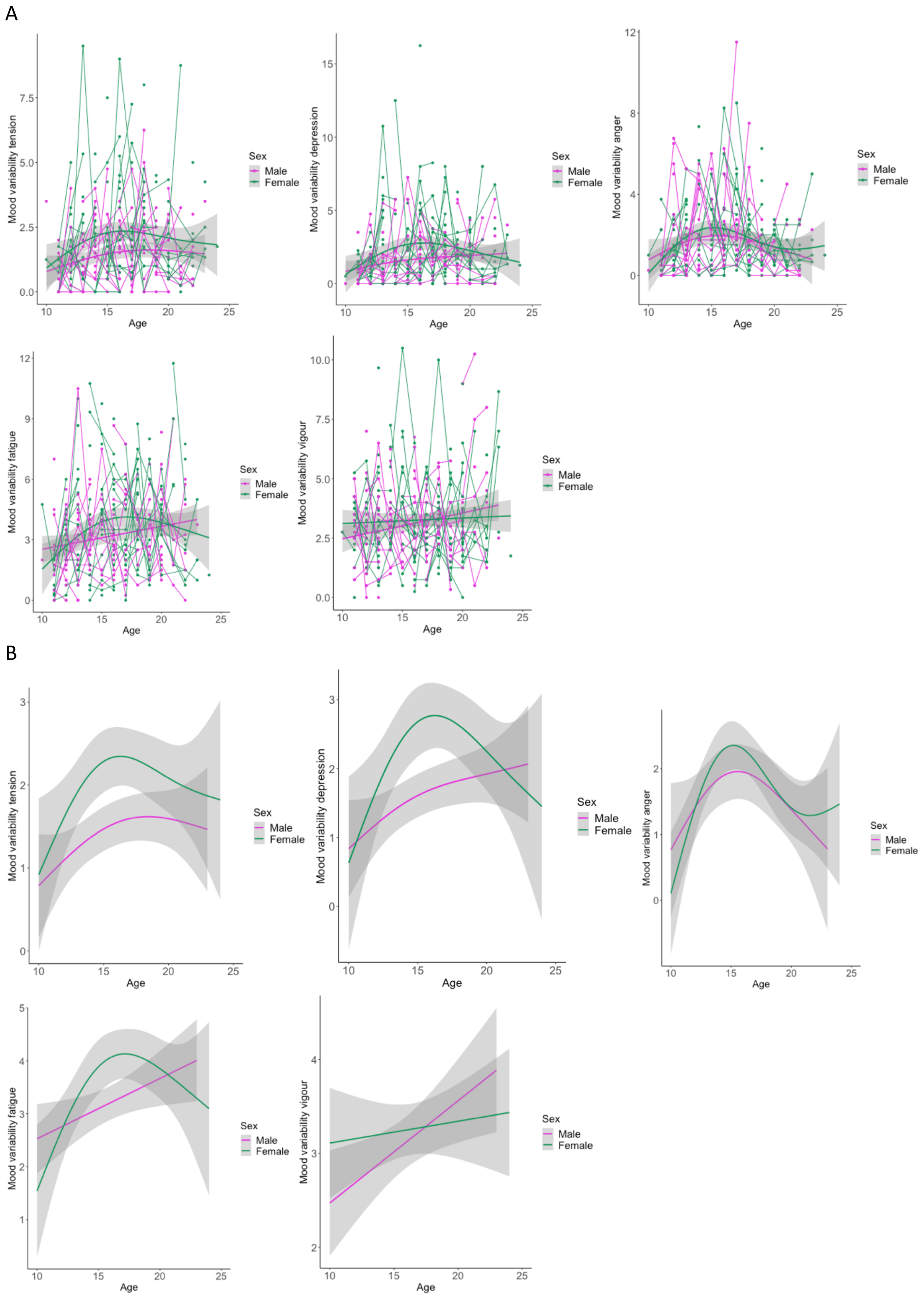


**Supplementary Figure S4. Development of mood variability per subscale (Tension, Depression, Anger, Fatigue and Vigor).** Plots with (A) and without individual trajectories (B).

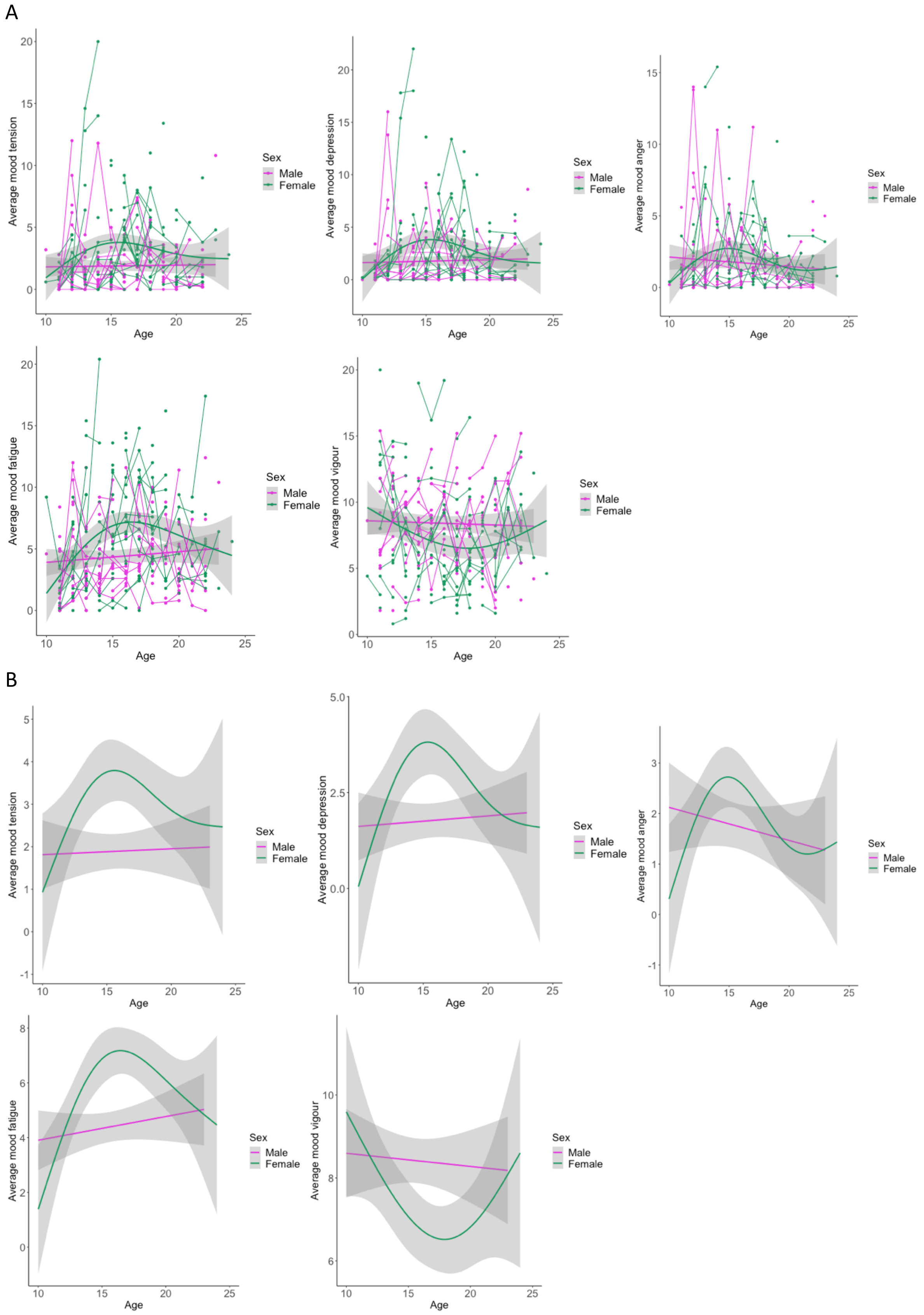


**Supplementary Figure S5. Development of average mood per subscale (Tension, Depression, Anger, Fatigue and Vigor).** Plots with (A) and without individual trajectories (B).

Since the development of the variability on the positive subscale, vigor, did not show an age with sex interaction effect, whereas the negative subscales did, a summary score consisting of the four negative subscales but not the positive subscale was used in the main analysis. This analysis was repeated with the 5 subscales and the results are reported below.

*Development of negative mood*
Mood variability and average mood (including vigor) showed a significant association with age by sex (BIC: 2431.39, p<0.001, $k$ = 4; BIC: 2426.60, p<0.001, $k$ = 4) in line with the results of the separate negative subscales (Supplementary Figure S6).

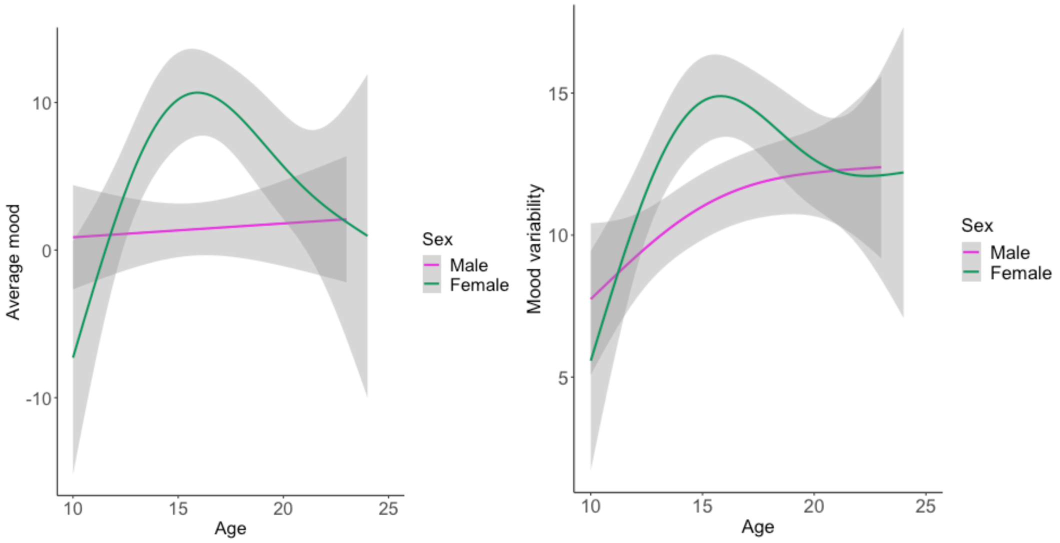


**Supplementary Figure S6. Development of average mood and mood variability**

*Association sleep and negative mood*
Similar to the main analysis, the association with objective and subjective sleep was examined. Sleep did not show an association with mood variability (Supplementary Table S3). Average mood did show an association with energy level, with higher energy levels compared to same-age peers, being associated with a more positive mood (Supplementary Figure S7).

**Supplementary Table S3. Association between sleep and mood**

| Sleep measure | Association with average mood ($p_{FDR}$) | BIC (k) | Association with mood variability ($p_{FDR}$) | BIC (k) |
|---|---|---|---|---|
| Objective sleep duration | 0.35 | 1151.11 (3) | 0.77 | 973.35 (3) |
| Sleep efficiency | 0.12 | 1147.10 (3) | 0.77 | 969.13 (3) |
| Subjective sleep duration | 0.89 | 2355.41 (3) | 0.85 | 1936.46 (3) |
| Energy level | <0.001 | 2483.68 (3) | 0.85 | 2054.97 (3) |

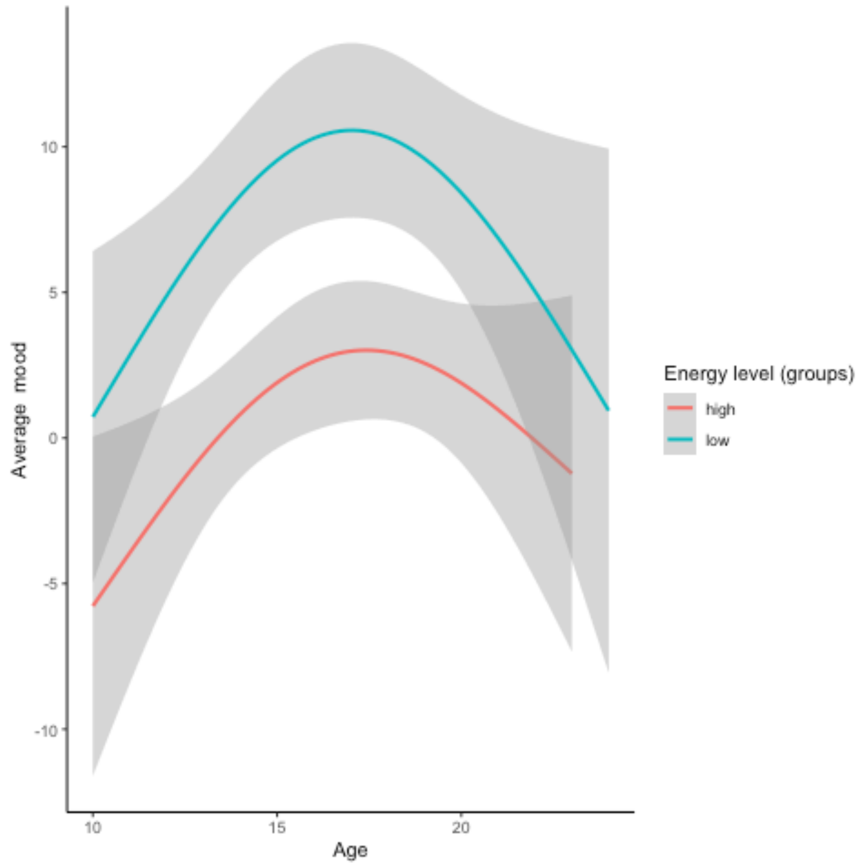


**Supplementary Figure S7. Association between average negative mood and energy level throughout development.**

*Association brain structure and negative mood*

Next, the association between mood variability and mood and brain structure throughout development was studied. The best-fit models showed that after correcting for average mood, mood variability showed an association with dlPFC on trend level (BIC: 2018.31, $p_{FDR}$=0.09, k=3) and vlPFC (BIC: 2021.15, $p_{FDR}$=0.09, k=3) thickness throughout development (Supplementary Figure S8). Those with thicker dlPFC and vlPFC in early and mid-adolescence, which are likely participants who are behind in development of their prefrontal cortex, show higher levels of mood variability. For dlPFC thickness this result was in line with the main analysis.

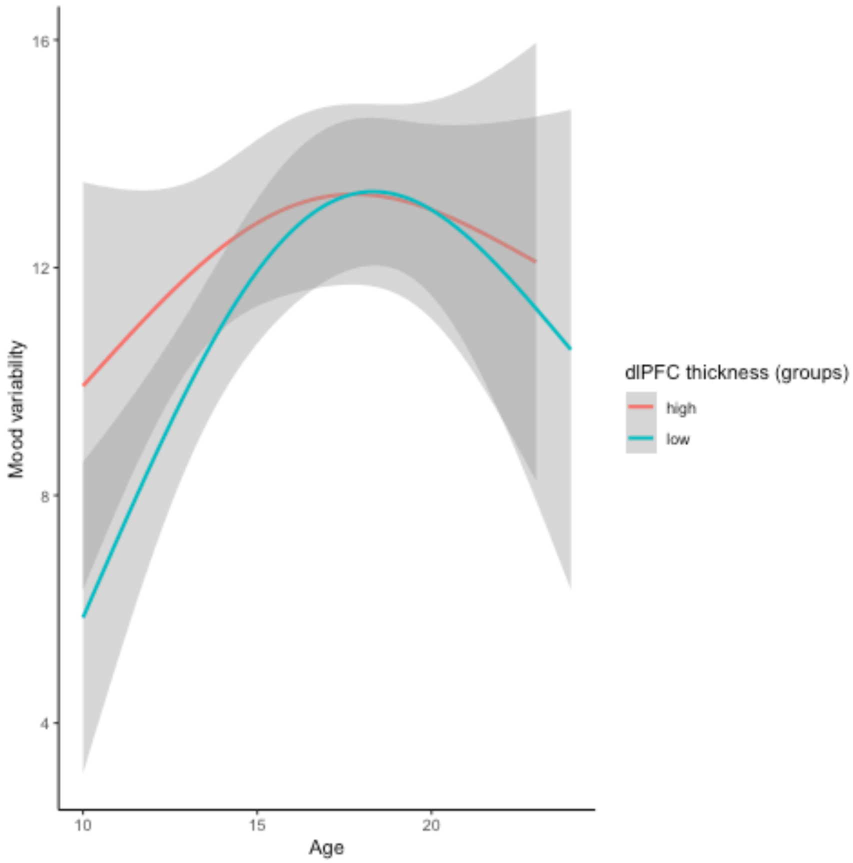


**Supplementary Figure S8. Association between mood variability and dlPFC thickness throughout development.**

### Supplementary Table S4. Correlation table

| | Negative mood variability | Average mood | Negative average mood | Anxiety symptoms | Depression symptoms | Subjective sleep duration | Subjective energy level | Objective sleep duration | Objective sleep efficiency | OFC thickness | ACC thickness | dlPFC thickness | vlPFC thickness | OFC surface area | ACC surface area | dlPFC surface area | vlPFC surface area | Ventral striatum | Amygdala |
|---|---|---|---|---|---|---|---|---|---|---|---|---|---|---|---|---|---|---|---|
| Mood variability | 0.968 | 0.524 | 0.520 | 0.346 | 0.364 | -0.289 | -0.210 | -0.164 | 0.095 | -0.155 | -0.077 | -0.166 | -0.161 | -0.050 | -0.001 | -0.081 | -0.207 | -0.105 | -0.081 |
| Negative mood variability | | 0.600 | 0.588 | 0.402 | 0.437 | -0.270 | -0.253 | -0.113 | 0.105 | -0.151 | -0.075 | -0.167 | -0.169 | -0.050 | -0.029 | -0.085 | -0.177 | -0.136 | -0.099 |
| Average mood | | | 0.970 | 0.510 | 0.629 | -0.244 | -0.337 | 0.022 | 0.172 | -0.186 | -0.054 | -0.147 | -0.184 | 0.012 | -0.048 | 0.000 | -0.075 | -0.174 | -0.142 |
| Negative average mood | | | | 0.472 | 0.578 | -0.247 | -0.267 | 0.013 | 0.174 | -0.148 | -0.065 | -0.102 | -0.120 | 0.050 | 0.003 | 0.069 | -0.062 | -0.116 | -0.047 |
| Anxiety symptoms | | | | | 0.735 | -0.151 | -0.247 | 0.054 | 0.068 | -0.066 | -0.001 | -0.027 | -0.056 | -0.222 | -0.094 | -0.191 | -0.092 | -0.204 | -0.219 |
| Depression symptoms | | | | | | -0.143 | -0.382 | -0.002 | 0.016 | -0.164 | -0.017 | -0.144 | -0.149 | -0.040 | -0.118 | -0.133 | -0.016 | -0.198 | -0.163 |
| Subjective sleep duration | | | | | | | 0.159 | 0.456 | -0.138 | 0.205 | 0.155 | 0.327 | 0.297 | 0.246 | 0.156 | 0.206 | 0.254 | 0.270 | 0.221 |
| Subjective energy level | | | | | | | | 0.017 | -0.005 | 0.139 | 0.009 | 0.215 | 0.231 | 0.033 | 0.164 | 0.099 | 0.075 | 0.153 | 0.216 |
| Objective sleep duration | | | | | | | | | 0.603 | 0.151 | -0.013 | 0.124 | 0.147 | 0.018 | 0.087 | 0.012 | 0.095 | 0.025 | 0.066 |
| Objective sleep efficiency | | | | | | | | | | 0.052 | -0.126 | -0.061 | 0.049 | -0.013 | 0.063 | -0.038 | -0.002 | -0.070 | -0.003 |
| OFC thickness | | | | | | | | | | | 0.483 | 0.715 | 0.793 | -0.051 | -0.073 | 0.213 | 0.132 | 0.385 | 0.166 |
| ACC thickness | | | | | | | | | | | | 0.443 | 0.389 | -0.023 | -0.260 | 0.102 | 0.104 | 0.128 | -0.030 |
| dlPFC thickness | | | | | | | | | | | | | 0.877 | 0.164 | 0.046 | 0.312 | 0.202 | 0.490 | 0.249 |
| vlPFC thickness | | | | | | | | | | | | | | 0.136 | 0.106 | 0.333 | 0.190 | 0.486 | 0.270 |
| OFC surface area | | | | | | | | | | | | | | | 0.636 | 0.776 | 0.679 | 0.622 | 0.554 |
| ACC surface area | | | | | | | | | | | | | | | | 0.639 | 0.443 | 0.525 | 0.422 |
| dlPFC surface area | | | | | | | | | | | | | | | | | 0.635 | 0.727 | 0.557 |
| vlPFC surface area | | | | | | | | | | | | | | | | | | 0.570 | 0.489 |

| | | | | | | | | | | | | | | | | | | | |
|---|---|---|---|---|---|---|---|---|---|---|---|---|---|---|---|---|---|---|---|
| Ventrale striatum | | | | | | | | | | | | | | | | | | | 0.664 |

**References**


1. Sadeh A (1989): Actigraphically based automatic bedtime sleep-wake scoring: validity and clinical application. *Journal of Ambulatory Monitoring* 2: 209–216.
2. Koopman-Verhoeff ME, Serdarevic F, Kocevska D, Bodrij FF, Mileva-Seitz VR, Reiss I, *et al.* (2019): Preschool family irregularity and the development of sleep problems in childhood: a longitudinal study. *Journal of Child Psychology and Psychiatry and Allied Disciplines* 60: 857–865.
3. Runze J, Euser S, Oosterman M, Dolan C v., Koopman-Verhoeff ME, Bakermans-Kranenburg MJ (2021): Actigraphic sleep and cortisol in middle childhood: A multivariate behavioral genetics model. *Comprehensive Psychoneuroendocrinology* 8: 100094.
4. Acebo C, Sadeh A, Seifer R, Wolfson AR, Hafer A, Carskadon MA (1999): Estimating Sleep Patterns with Activity Monitoring in Children and Adolescents: How Many Nights Are Necessary for Reliable Measures? *SLEEP*, vol. 22. Retrieved from https://academic.oup.com/sleep/article/22/1/95/2731704
5. Hansen M, Janssen I, Schiff A, Zee PC, Dubocovich ML (2005): The impact of school daily schedule on adolescent sleep. *Pediatrics* 115: 1555–1561.
6. DeMasi O, Feygin S, Dembo A, Aguilera A, Recht B (2017): Well-being tracking via smartphone-measured activity and sleep: Cohort study. *JMIR Mhealth Uhealth* 5. https://doi.org/10.2196/mhealth.7820
7. Harezlak J, Ryan LM, Giedd JN, Lange N (2005): Individual and population penalized regression splines for accelerated longitudinal designs. *Biometrics* 61: 1037–1048.
